# Genome-wide analysis of mRNA decay in Arabidopsis shoot and root reveals the importance of co-translational mRNA decay in the general mRNA turnover

**DOI:** 10.1101/2024.03.23.586374

**Authors:** Carpentier Marie-Christine, Receveur Anne-Elodie, Boubegtitene Alexandre, Cadoudal Adrien, Bousquet-Antonelli Cécile, Merret Rémy

**Affiliations:** CNRS-LGDP UMR 5096, 58 avenue Paul Alduy 66860 Perpignan, France; Université de Perpignan Via Domitia, LGDP-UMR5096, 58 avenue Paul Alduy, 66860 Perpignan, France

## Abstract

Until recently, the general 5’-3’ mRNA decay was placed in the cytosol after the mRNA was released from ribosomes. However, the discovery of an additional 5’ to 3’ pathway, the Co-Translational mRNA Decay (CTRD), changed this paradigm. Up to date, defining the real contribution of CTRD in the general mRNA turnover has been hardly possible as the enzyme involved in this pathway is also involved in cytosolic decay. Here we overcame this obstacle and created an Arabidopsis line specifically impaired for CTRD called XRN4ΔCTRD. Through a genome-wide analysis of mRNA decay rate in shoot and root, we tested the importance of CTRD in mRNA turnover. First, we observed that mRNAs tend to be more stable in root than in shoot. Next, using XRN4ΔCTRD line, we demonstrated that CTRD is a major determinant in mRNA turnover. In shoot, the absence of CTRD leads to the stabilization of thousands of transcripts while in root its absence is highly compensated resulting in faster decay rates. We demonstrated that this faster decay rate is partially due to the XRN4-dependent cytosolic decay. Finally, we correlated this organ-specific effect with XRN4ΔCTRD line phenotypes revealing a crucial role of CTRD in mRNA homeostasis and proper organ development.

## INTRODUCTION

The tight control and dynamic fine-tuning of gene expression are *sine qua non* conditions for life necessary to maintain proper cellular homeostasis and identity in response to developmental and environmental cues. The reprograming of gene expression is exerted at many steps: transcriptionally, post-transcriptionally at the level of messenger RNA (mRNA) and post-translationally at the protein level. All these processes are studied intensively in a global effort to understand basic mechanisms regulating gene expression in eukaryotes. Regulations at the mRNA level emerged as potent means to modify gene expression pattern to swiftly adapt the cellular activity and allow organisms to acclimate to and survive stress (1).

In the cytoplasm, over its entire lifetime, any mRNA is in balance between translation, storage and decay. The spatiotemporal regulation of this equilibrium is critical for transcriptome adjustment in response to developmental and environmental cues in plants (2). In *Arabidopsis thaliana*, alterations in mRNA decay due to the loss of function of key factors often result in post-embryonic lethality, severe growth defects or impairment of the stress response (3–5). As an example, the loss of function of XRN4, the major cytoplasmic 5’-3’ exoribonuclease, induces defects in leaf morphology, root growth and stress response such as dark response, nitrogen deprivation or heat stress (6–9). These pleiotropic phenotypes strongly support the importance of mRNA decay across development and response to stress in plants.

Degradation of mRNAs is an important component of gene expression that controls the steady state concentration of functional transcripts in the cell. This process is well conserved in eukaryotes and can initiate from the 5’ or 3’ extremities of mRNA (10). The so-called general 5ʹ to 3ʹ mRNA turnover that takes place in the cytoplasm occurs along three steps. Removal of the poly(A) tail of mRNA by deadenylases is the first and rate-limiting step in mRNA degradation. Following poly(A) tail shortening (deadenylation), the decapping complex (composed of VCS, DCP1 and DCP2 in *Arabidopsis thaliana*) hydrolyses the mRNA cap structure. Then, the 5ʹ-phosphate end of the decapped mRNA is attacked by the cytoplasmic exoribonuclease (XRN4 in *Arabidopsis thaliana*, XRN1 in *Saccharomyces cerevisiae*) which digests the body of the transcript. The 3’ to 5’ mRNA decay can act through the activity of the RNA exosome complex or the suppressor of varicose, SOV in *Arabidopsis thaliana* (11, 12). The biological relevance of these pathways was addressed in *Arabidopsis thaliana*, revealing a large contribution of 5’-3’ mRNA decay in the general mRNA turnover (13). A defect in one of these pathways can be compensated by a feedback mechanism to maintain proper mRNA homeostasis. However, the molecular mechanism governing this feedback is still unclear (10, 13). The redundancy of the 5’-3’ and 3’-5’ decay pathways was also demonstrated. Indeed, dysfunction of bidirectional RNA decay pathways results in an accumulation of spurious siRNAs and causes several developmental defects (14), demonstrating the importance of mRNA decay in plant development.

While initially thought to be mutually exclusive, data from several organisms including *Saccharomyces cerevisiae* and *Arabidopsis thaliana* now proved that mRNA decay and translation are actually tightly intertwined (15–17). In particular, the existence of an evolutionarily conserved turnover process, the 5’-3’ co-translational mRNA decay (CTRD) was proposed, where mRNAs are degraded from 5’ to 3’ while still engaged in translation. The existence of a CTRD was first proposed in yeast (18). The authors demonstrated using a set of reporter genes that uncapped mRNAs can be detected associated with ribosomes. Next the existence of this pathway in *Arabidopsis thaliana* was also proposed (7, 19). Heat stress induces a slowing down of ribosome elongation triggering the CTRD of transcripts coding for proteins with hydrophobic N-termini (19). This process was subsequently found to globally shape the whole transcriptome of many eukaryotes such as yeast, mammalian cells and plants (15–17, 20, 21). Recently, this pathway was proposed to be conserved in Angiosperms (21).

This transcriptome-wide effect of CTRD was revealed by sequencing of RNA decay intermediates using high-throughput degradome (or 5’P-Seq) approaches (20, 22–24). These approaches revealed that mRNA decay intermediates follow an XRN1/4-dependent, three-nucleotide periodicity. This periodicity can be explained by the fact that XRN1/4 chases the last translating ribosome in a codon-by-codon manner, and since it is a processive enzyme, only degradation intermediates protected by ribosomes can be captured. Consequently, each of these degradome approaches gives a snapshot of CTRD and also reveals ribosome dynamics and how degradation impacts this dynamic (15). Since the development of 5’P-Seq approaches an increasing number of articles reported the existence of CTRD, identified actors of this process and provided evidence that CTRD activity is variable across conditions (16, 20–22, 25–27). However, the actual respective contributions of the 5’-3’ cytosolic and 5’-3’ CTRD decays to the general 5’-3’ mRNA turnover have never been properly measured in any organisms.

Until now, it has been difficult to distinguish the relative contributions of 5’-3’ cytosolic and co-translational decays in mRNA turnover since both processes involve the same enzyme: XRN1/XRN4. Here, we overcame this obstacle and created an Arabidopsis line specifically impaired for CTRD called XRN4ΔCTRD line. In this line, XRN4 can still decay mRNAs in the cytosol but can no longer perform CTRD. Using this line in comparison to WT and *xrn4* mutant, we performed a genome-wide mRNA decay analysis respectively in shoot and root. First, we demonstrated that mRNAs expressed in both organs tend to be more stable in root than in shoot. Next, we demonstrated that CTRD is the main 5’-3’ mRNA decay pathway in shoot. Indeed, 70% of transcripts targeted by 5’-3’ mRNA decay are targeted by CTRD with most of them exclusively targeted by this pathway. Finally, in root, we observed that an RNA decay feedback mechanism takes place specifically in the absence of CTRD, resulting in increased decay rates. We showed that part of this feedback is mediated by the XRN4-dependent cytosolic decay. This mechanism induces specific root phenotypes in the XRN4ΔCTRD line opposite to those observed in the *xrn4* mutant.

## MATERIAL AND METHODS

### Sequence analysis

Sequence alignment was performed with Uniprot protein sequence of ScXRN1 (P22147), CeXRN1 (Q9BHK7), HsXRN1 (Q8IZH2), ScXRN2 (Q02792), AtXRN4 (Q9FQ04), HsXRN2 (Q9H0D6), CeXRN2 (Q9U299) and DmXRN1 (Q9VWI1).

### 3D structure prediction

3D structure prediction was performed using Alphafold software (28–30). PDB files corresponding to AtXRN4 (Q9FQ04) and ScXRN1 (P22147) were loaded simultaneously on Alphafold 3D viewer to compare 3D prediction models using default parameters. Regions corresponding to L1, L2 and L3 loops in ScXRN1 (Positions 46 to 244) were then selected to generate Figure 1B.

**Figure 1.**
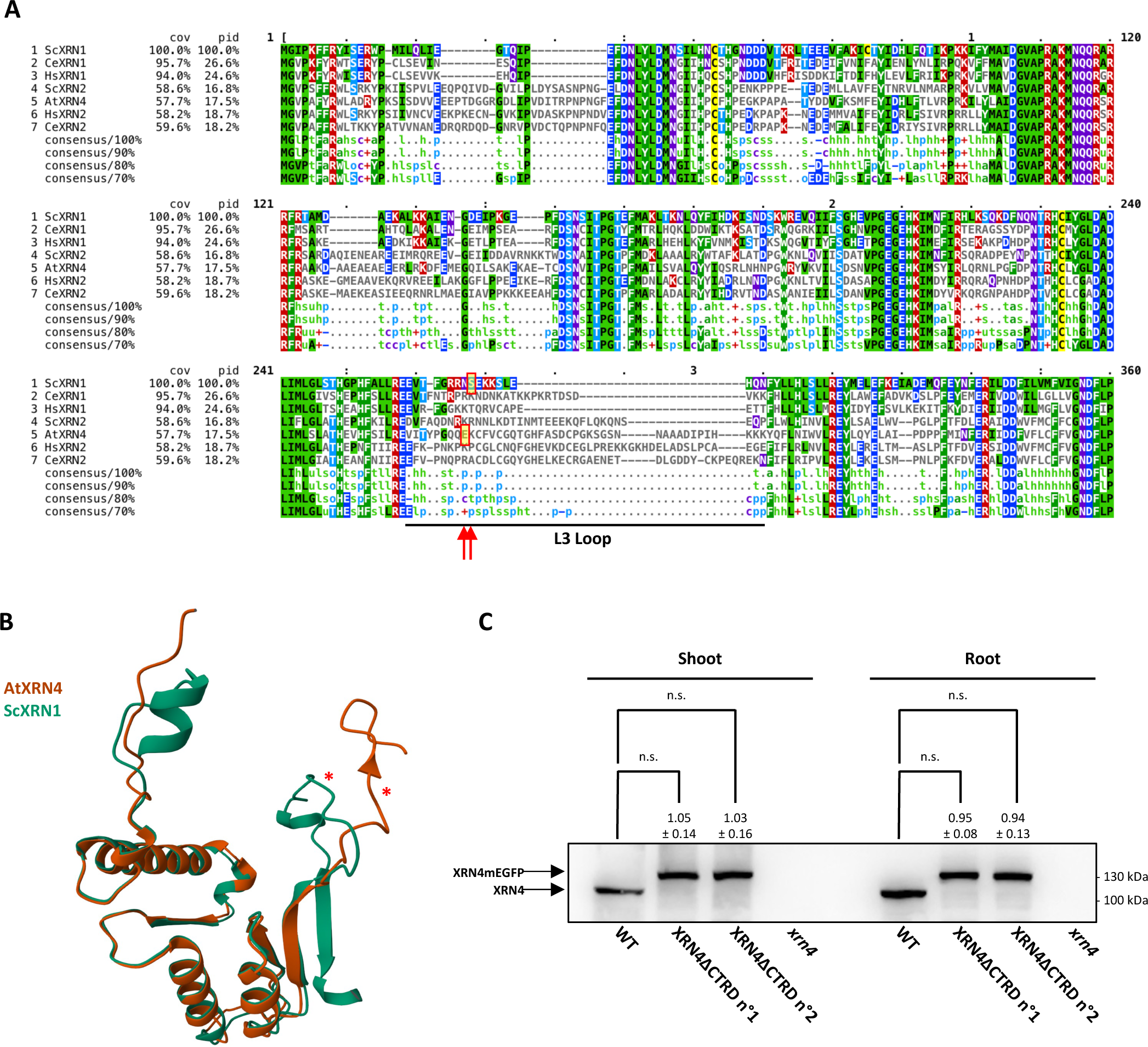
Generation of the AtXRN4ΔCTRD transgenic line. A. Alignment of the *Arabidopsis thaliana* XRN4 protein with orthologs from Ce, *Caenorhabditis elegans*; Hs, *Homo sapiens*; Sc, *Saccharomyces cerevisiae*. Only the first 360 amino acids are only represented. The position of the L3 loop in *Saccharomyces cerevisiae* is presented (positions 227-244). To generate the AtXRN4ΔCTRD line, the mEGFP was inserted after the position E263. Red arrows indicate positions of S235 in ScXRN1 and E263 in AtXRN4. B. Superposition of ScXRN1 and AtXRN4 AlphaFold prediction. Regions corresponding to L1, L2 and L3 loops in ScXRN1 are only represented. Red asterisks indicate the location of mEGFP insertion. C. XRN4 accumulation in XRN4ΔCTRD lines compared to Wild-type (Col0) and *xrn4* using XRN4 specific antibody. The fold change relative to Wild-type is indicated. N= 4 biological replicates. A t-test was performed to test significance. n.s., not significant.

### Generation of the AtXRN4ΔCTRD and AtXRN4ΔCTRD(R_118_A/R_119_A) transgenic lines

The mEGFP coding sequence was inserted in frame after the triplet encoding for E263 within the XRN4 CDS sequence (Supplemental Figure 1). This sequence was then subcloned in a gateway vector containing XRN4 promoter sequence (1100 bp before XRN4 start codon). XRN4ΔCTRD was transformed in *xrn4-5* (SAIL_ 681_E01) background using the *Agrobacterium tumefaciens* based floral dip technique (31). Independent lines expressing XRN4ΔCTRD were screened by western-blot using XRN4 specific antibody (7). Two independent lines expressing XRN4 at levels similar to WT were selected for further analysis. To generate AtXRN4ΔCTRD(R_118_A/R_119_A) transgenic lines, the same procedure was followed, except that Arg118 and Arg119 were mutated in Alanine prior to cloning steps.

### Growth conditions

Analyses were carried out with Columbia-0 line as Wild-type (WT), XRN4ΔCTRD line and *xrn4-5* mutant (SAIL_681_E01). Seeds were sown on a 245×245mm square plate in a single row. Plantlets were grown vertically on synthetic Murashige and Skoog medium (Duchefa) containing 1% Sucrose and 0.8% plant agar at 22°C under a 16-h-light/8-h-dark regime.

### Confocal microscopy analysis

Confocal microscopy was performed on 7-d-old seedlings. Prior to confocal microscopy analysis, seedlings were fixed using 2% formaldehyde. After DAPI staining, slides were mounted in Vectashield. Observations and acquisitions were performed using LSM700 (Zeiss) confocal microscope.

### Transcriptional arrest time course

Transcriptional inhibition was performed on 15-d-old seedlings as described previously (13). The time-course experiment was performed 2 hours after daybreak. Plantlets were transferred horizontally in an incubation buffer (15 mM sucrose, 1 mM Pipes pH 6.25, 1 mM KCl, 1 mM sodium citrate, 1 mM cordycepin) in a single row to easily separate shoots and roots. Roots and shoots were collected separately at 0, 7.5, 15, 30, 60, 120, 240 and 480 min after transcription arrest. Roots and shoots were separated using a scissor at the basis of the hypocotyl and rapidly transferred to liquid nitrogen prior to RNA extraction. For RNAseq, four biological replicates per genotype were collected.

### Total RNA extraction, RNA sequencing

Total RNA was isolated using Monarch Total RNA Miniprep Kit according to manufacturer’s instructions. Total RNA was then subjected to ribosomal RNA depletion using QIAseq FastSelect rRNA Plant Kit according to manufacturer’s instructions. RNA library preparation was performed using QIAseq Stranded RNA Library Kit. The samples were multiplexed and sequenced in PE 2×150. At least 20 million of clusters were obtained per sample. After filtering out reads corresponding to chloroplastic, mitochondrial, ribosomal, and small RNA sequences, reads were mapped against the TAIR10 genome using Hisat2 v2.03 and the gtf Araport11 annotation file with standard parameters. Read counts by gene were performed by htseq-count in RPM. Only transcripts with at least 1 RPM at T_0_ in all genotypes and all replicates were kept. Differential expression analyses were performed using DESeq2.

### mRNA decay rate analysis

mRNA decay rate analysis was performed according to (13). Data normalization, as well as the modelling of mRNA decay and genotype effect, were performed using the Bioconductor RNAdecay package (13, 32). Data normalization was performed using the mean fold increase of 50 stable genes in all genotypes (Supplemental Table 1). mRNA half-life (t_1/2_) for each transcript in each genotype is presented on Supplemental Table 2.

### Polysome profile analysis

Polysome profiles and western-blot analysis were performed as described previously (16). To quantify XRN4 association with ribosomes, fractions corresponding to monosome and polysomes were pooled. To quantify free XRN4, fractions corresponding to free mRNPs were pooled. Proteins were then precipitated from both fractions by adding 2 volumes of absolute ethanol. After 6 hours at 4°C and centrifugation, protein pellets were washed 5 times with ethanol 70%. Finally, pellets were resuspended in Laemmli 4X buffer. For western-blot analysis, the same percentage of each fraction (Input, free mRNP and ribosome-bound fractions) were loaded. Quantification was performed using Vilber software on 4 biological replicates. All blots were prepared and immunoblotted in parallel and simultaneously exposed for chemiluminescence quantification.

### Immunoblotting

XRN4 antibody (7) was used at 1/1000th dilution. UGPase antibody was purchased (Agrisera) and used at 1/5000th dilution. Antibodies against DCP5, PAT1, PAT1H1, PAT1H2 were produced in rabbits using the Eurogentec double X immunization program. The DCP5 protein was detected using the serum affinity-purified against peptide PNGHSFPNHNGYRGRG at 1/1000th dilution; the PAT1 protein was detected using the serum affinity-purified against peptide VEQRIPDRTKLYPEP at 1/1000th dilution; the PAT1H1 protein was detected using the serum affinity-purified against peptide PGNRSPQASPGNLHR at 1/1000th dilution; the PAT1H2 protein was detected using the serum affinity-purified against peptide VPPRVSNHGNPNDGL at 1/1000th dilution. Primary antibody was incubated overnight at 4°C under constant agitation. A horseradish peroxidase-coupled antibody was used as secondary antibody. Signal was revealed with the Immobilon-P kit from Millipore.

### RT-ddPCR

A spike-in Luciferase RNA was first transcribed *in vitro* using HiScribe™ T7 Quick High Yield RNA Synthesis Kit (New England Biolabs) according to Manufacturer’s instructions. The FLuc plasmid provided in the kit was used as template. Luciferase RNA was then purified using Monarch RNA cleanup kit (New England Biolabs) according to Manufacturer’s instructions. For each sample, 500 ng of total RNA was spiked with 10 pg of Luciferase RNA and reverse-transcribed using SuperScript IV kit using random primers (Thermo Scientific). cDNAs were then diluted 50-fold prior to ddPCR analysis. ddPCR was performed as described previously (33). Luciferase quantification in each sample was used for normalization. Primers used for ddPCR are listed on Supplemental Table 3.

### 5’P sequencing

5’Pseq library was prepared as described previously (34). Raw reads were trimmed to 50 pb before mapping. Metagene analysis was performed using FIVEPSEQ software v1.0.0 (22). “meta_counts_START.txt” and “meta_counts_STOP.txt” files were used to analyse 5’P reads accumulation around start and stop positions. The translational termination stalling index (TSI) was defined as the ratio of the number of 5’P read ends at the ribosome boundary (16–17 nt upstream from stop codon) to the mean number of 5’P read ends within the flanking 100 nt. Transcripts with a TSI value higher than 3 in WT were used to assess CTRD activity in *xrn4* and XRN4ΔCTRD lines.

### GO terms analysis

GO terms analysis was performed using clusterProfiler R package.

### Phenotypic analysis

For leaf phenotypes, plants were cultivated in soil in a growth chamber under long day conditions (16h light at 21°C/8h dark at 16°C) at 65% relative humidity. Number of leaves and leaf fresh weight were calculated on 34 days-old plants. For root phenotypes, plants were cultivated *in vitro* on synthetic MS containing 1% Sucrose and 0.8% plant agar at 22°C under a 16-h-light/8-h-dark regime. For NaCl treatment, 4-d-old seedlings were transferred to MS media supplemented with 125 mM NaCl. Root length was then determined on 15-d-old seedlings.

## RESULTS

### Generation of the AtXRN4ΔCTRD transgenic line

As XRN4 catalyses both cytosolic and co-translational mRNA decay, it is challenging to uncouple these pathways and determine their respective contributions. To address this question, we took advantage of a recent structural analysis of the yeast ScXRN1 physical interactions with the 80S ribosome (35). ScXRN1 carries several ribosome contact areas, including three loops (L1, L2 and L3), with the flexible loop L3 located in between the CR1 and CR2 conserved structured regions that form the catalytic domain, that is essential for ribosome binding (Figure 1A) (35). Tessina *et al.*, showed that disruption of loop L3, through insertion of a monomeric GFP (mEGFP) abolishes ScXRN1 association with polysomes without affecting its exoribonucleolytic activity. We reasoned that in such mutant co-translational mRNA decay should be impaired. As we are interested in understanding the molecular mechanisms of 5’-3’ mRNA degradation in *Arabidopsis thaliana,* we set to construct a line solely expressing an *XRN4* allele unable to bind to polysomes. As we and others previously showed (7), the first half of XRNs, encompassing the CR1-CR2 catalytic core is highly conserved at the primary sequence level, and an AlphaFold-based 3D structure prediction supports the existence of loop L3, at the end of the CR1 in AtXRN4 as in ScXRN1 (Figure 1B). In yeast, the mEGFP was inserted into loop L3, after serine 235, nine amino acids after the conserved CR1 region (Figure 1A), we hence performed a similar approach in AtXRN4 by inserting mEGFP after glutamic acid 263 (AtXRN4 numbering) (Figure 1A). We next generated a stable transgenic line expressing this modified *XRN4* under the control of its own promoter in an *xrn4* loss-of-function mutant (*xrn4-5*, SAIL_681_E01). We called this line XRN4ΔCTRD and identified two independent transgenic plants expressing XRN4ΔCTRD at levels similar to that of the endogenous protein in both shoot and root (Figure 1C). Several approaches were then deployed to demonstrate that this line is impaired in co-translational mRNA decay (Figures 2-3 and Supplemental Figure 2). First, we quantified XRN4 association with ribosomes by polysome fractionation followed by western-blotting using XRN4 specific antibodies. Fractions corresponding to monosomes and polysomes were pooled and referred to as “Ribosome-bound fractions”. Fractions corresponding to free mRNPs were also pooled and used to quantify free XRN4. XRN4 signal was compared in both fractions relative to an input fraction prior to polysome fractionation (Figure 2A-B). Interestingly, in both shoot and root, XRN4 signal is significantly higher in the ribosome-bound fractions than in the free mRNP fractions suggesting that CTRD activity is important in both organs. We performed a similar analysis with the XRN4ΔCTRD line and found that XRN4 signal in ribosome-bound fractions drastically decreased (Figure 2A-B). In contrast, a GFP insertion at the C-terminus of XRN4 does not affect XRN4 distribution (Figure 2A-B).

**Figure 2.**
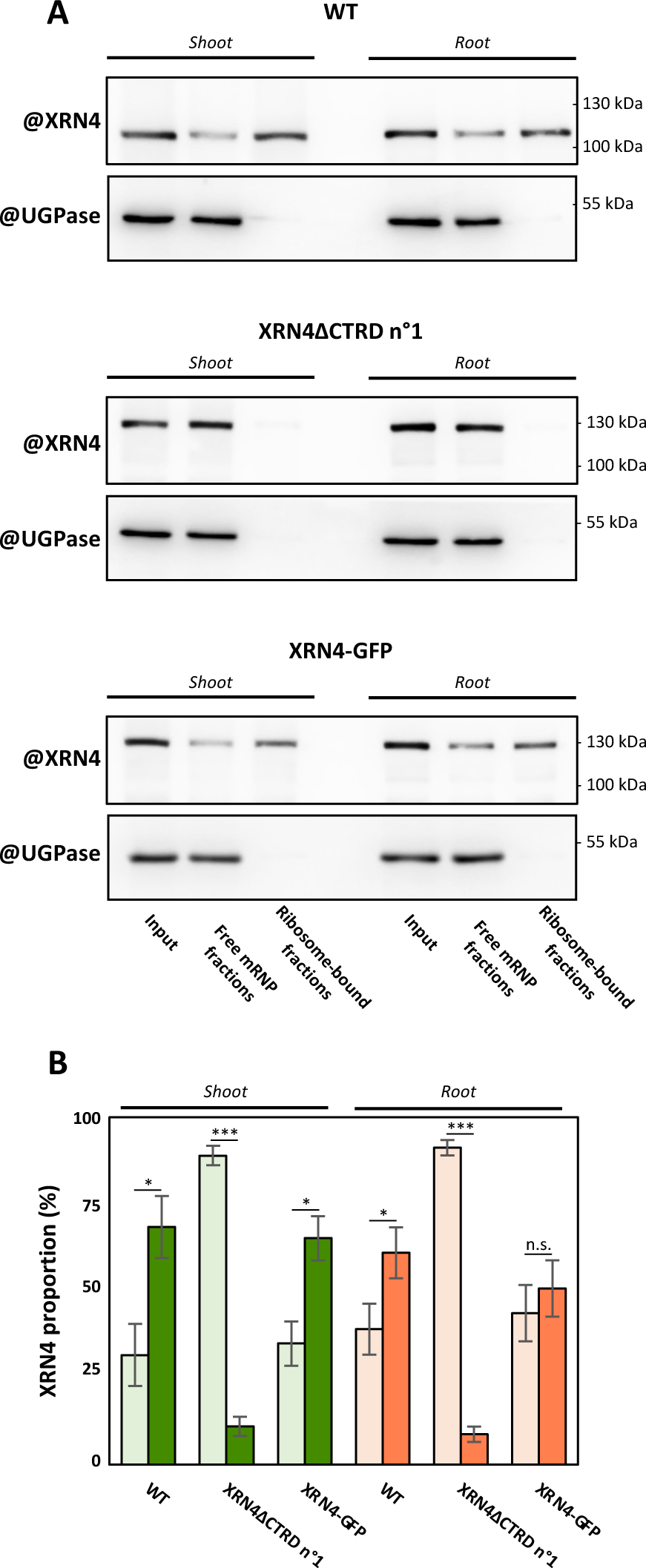
XRN4 association to polysomes is impaired in AtXRN4CTRD transgenic line. A. Distribution of XRN4 in free mRNP and ribosome-bound fractions in WT, XRN4ΔCTRD line and XRN4-GFP complemented line using XRN4 specific antibody. The same percentage of each fraction was loaded. UGPase antibody was used as a negative control. B. Proportion of XRN4 in free mRNP (light colored bar) and ribosome-bound (dark colored bar) fractions in each line. N= 4 biological replicates. A t-test was performed to test significance. All blots were prepared and immunoblotted in parallel and simultaneously exposed for chemiluminescence quantification.

**Figure 3.**
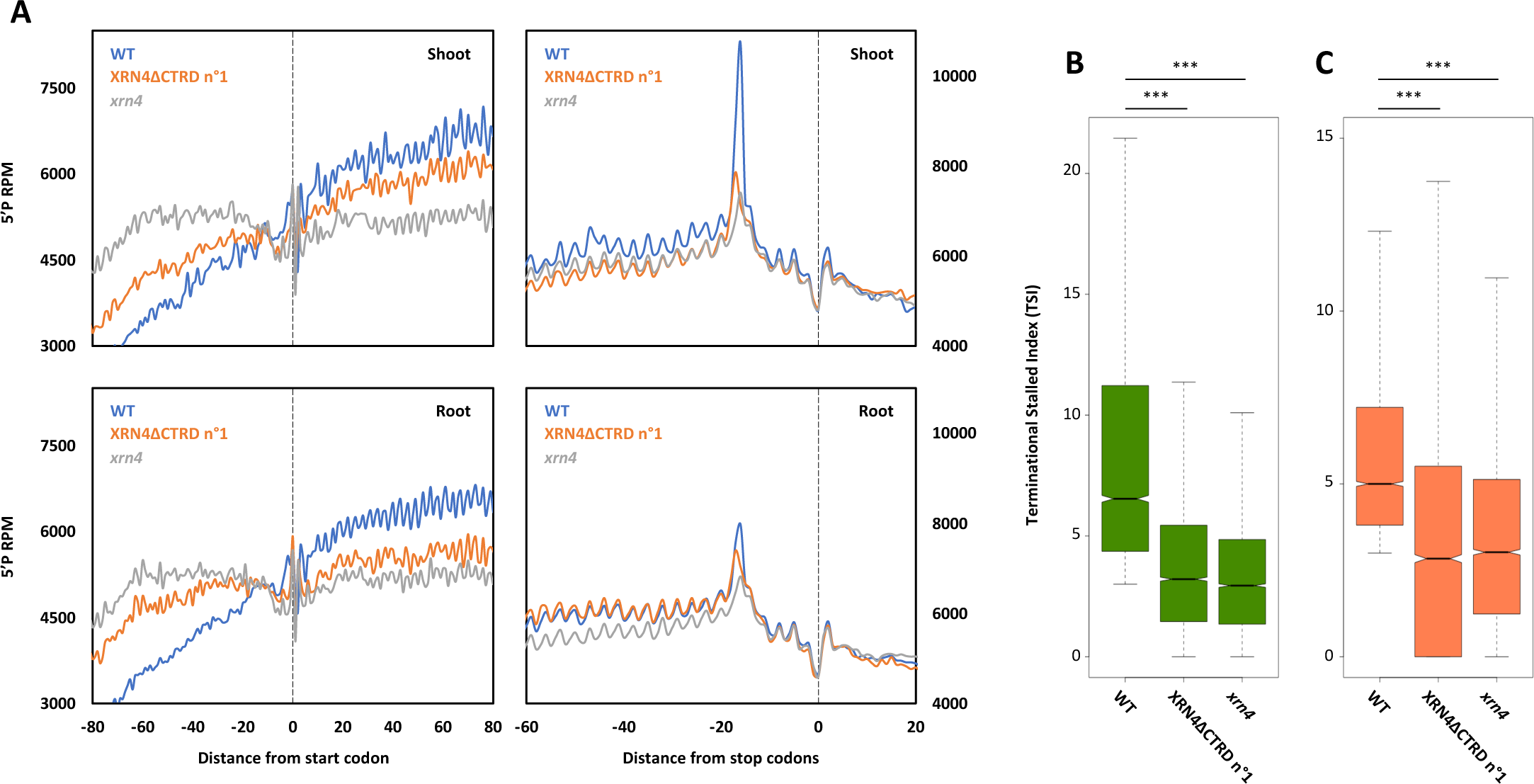
AtXRN4CTRD transgenic line is impaired in co-translational mRNA decay. A. Metagene analysis of 5’P reads accumulation around start and stop codons in both shoot and root. B, C. Distribution of Terminational Stalled index (TSI) in both shoot and root. N= 2 biological replicates. A Wilcoxon test was performed to test significance.

Then, to test CTRD activity in XRN4ΔCTRD lines, we ran 5’Pseq assays and compared results to that of wild-type (WT) and *xrn4* mutant through monitoring of reads accumulation around stop and start codons (Figure 3A). As expected, a meta-transcriptome analysis shows a strong overaccumulation of reads at position 16/17nt before the stop codon in wild-type, a hallmark of active CTRD (16). In the XRN4ΔCTRD line, the minus 16/17nt peak shows a dramatic decrease that is almost identical to that of the *xrn4* mutant, supporting a drastic inactivation of CTRD (Figure 3A). Conversely, before the start codon, a clear over-accumulation (as compared to WT) of 5’P reads is observed in both *xrn4* and XRN4ΔCTRD lines (Figure 3A), another hallmark of 5’-3’ mRNA decay impairment. We suppose that the higher accumulation observed in *xrn4 c*ompared to XRN4ΔCTRD line is the consequence of the loss of both the cytosolic and co-translational mRNA decays. Recently, it was reported that the terminational stalling index (TSI) can be used to identify CTRD activity at individual transcript levels (21, 36). Using this TSI metric, we demonstrated that CTRD activity is reduced in the XRN4ΔCTRD line to levels similar to that of a complete loss of XRN4 (Figure 3B-C). Finally, we observed through confocal microscopy that the XRN4ΔCTRD shows a subcellular distribution similar to that of a C-terminally tagged XRN4-GFP fusion (Supplemental Figure 2). In both lines, XRN4 was found to be distributed in the cytoplasm with few cytoplasmic foci as previously described (7, 37). All together, these data support that the insertion of a monomeric GFP into AtXRN4 loop 3, does not affect the total levels of the protein nor does it impair its cytoplasmic accumulation. But that it blocks its ability to associate to polysomes, and inactivates the CTRD pathway as efficiently as total loss of the XRN4 protein. We hence conclude that the XRN4ΔCTRD lines only retains the ability to degrade mRNAs outside of polysomes, providing a useful tool to evaluate the contribution of CTRD to the 5’-3’ turnover of transcripts.

### Strategy for mRNA half-life determination and CTRD contribution

To determine the contribution of CTRD in mRNA turnover, we took advantage of the transgenic line XRN4ΔCTRD. A genome-wide mRNA decay analysis was carried on WT (Col0, active cytosolic and CTRD degradations), *xrn4* mutant (*xrn4-5*, SAIL_681_E01, inactive cytosolic and CTRD degradations) and two independent XRN4ΔCTRD lines (active cytosolic and inactive CTRD degradations) 15-d-old seedlings as we have shown that CTRD is highly active at this stage (16). As little is known about the importance of CTRD at the organ level, the analysis was performed on root and shoot separately (Supplemental Figure 3A). After transcriptional inhibition by addition of cordycepin, roots and shoots were collected separately at 0, 7.5, 15, 30, 60, 120, 240 and 480 min. In order to capture both deadenylated and adenylated mRNA decay intermediates, rRNA depletion was performed prior to RNA library preparation. RNA libraries were then multiplexed and sequenced in Paired-Ends 2X150bp. RNAseq was performed on four biological samples, revealing reproducible differences between genotypes and time-points (Supplemental Figure 4). Only transcripts with at least 1 RPM at T_0_ in all genotypes and all replicates were kept. Prior to mRNA decay analysis, a DESeq analysis was performed between the two XRN4ΔCTRD lines (Supplemental Figure 5). Among the 16 comparisons, only 782 unique transcripts were detected as significantly up- or down-regulated demonstrating the reproducibility of both lines. These transcripts were removed from the analysis. Finally, to simplify subsequent analyses, the mean of RPM values of both XRN4ΔCTRD lines was performed resulting in one XRN4ΔCTRD line. The decay analysis was thus initiated on 12 306 transcripts (Supplemental Figure 3B). To determine the mRNA half-life for each transcript, we took advantage of the development of an RNA decay pipeline based on a mathematical modeling approach (13, 32). After data normalization and mathematical modelling, an mRNA half-life (t_1/2_) was obtained for the 12 306 transcripts in WT, *xrn4* and XRN4ΔCTRD lines in both root and shoot (Supplemental Table 2).

### mRNA decay landscape between shoot and root in WT

The study was first initiated in WT comparing shoot and root data. The analysis of mRNA decay rates in both organs reveals a wide range of decay from very fast (few minutes) to very slow (several hours). Interestingly, mRNAs expressed in both organs tend to be more stable in root than in shoot (171 minutes versus 103 minutes) (Figure 4A-B, Supplemental Table 2). The mRNA half-life distribution shows that root presents less unstable (t_1/2_ < 100 minutes) and more stable mRNAs (t_1/2_ > 100 minutes) compared to shoot. We next tested if mRNA stability in each organ could be correlated with corresponding biological functions. For each transcript, the fold change between its half-life in shoot versus root (FC t_1/2 shoot /_ t_1/2 root_) was determined and followed by a GO slim analysis (Figure 4C). Interestingly, some biological functions present a higher stability in shoot than in root and vice-versa. mRNAs stability in *Arabidopsis thaliana* was correlated with some *cis*-elements such as intron numbers and 5’UTRs A/G content (13). We analyzed the distribution of these two features and also observed a strong correlation between intron number, 5’UTR content and mRNA stability at the organ level (Supplemental Figure 6). Intron number, 5’UTR increased A content and 5’UTR decreased G content were positively correlated with mRNA stability both in shoot and root. As median mRNA half-life is much higher in root than in shoot, we then tested if this difference could be due to different levels of key components of the 5’-3’ and 3’-5’ mRNA decay machineries. Interestingly, the *SKI2* mRNA (an RNA helicase, subunit of the SKI complex involved in the 3’-5’ decay process) is much more stable in shoot than in root (787 minutes in shoot and 218 minutes in root) (confirmed by an independent approach, Supplemental Figure 7), suggesting that the 3’-5’ mRNA decay could be less active in root than in shoot. Regarding the 5’-3’ mRNA decay components, we analyzed published quantitative proteomic data and found that PAT1H1 and PAT1H2 (cofactors of decapping) are less abundant in root compared to shoot (38). Using specific antibodies against PAT1, PAT1H1 and PATH2, we tested protein accumulation in our conditions (Supplemental Figure 8). While XRN4 and DCP5 seemingly accumulate at similar levels in root and shoot, we found that PAT1, PAT1H1 and PAT1H2 proteins under accumulate in root compared to shoot. These data could suggest that co-factors of decapping are less abundant in root resulting in a lower 5’-3’ decay activity.

**Figure 4.**
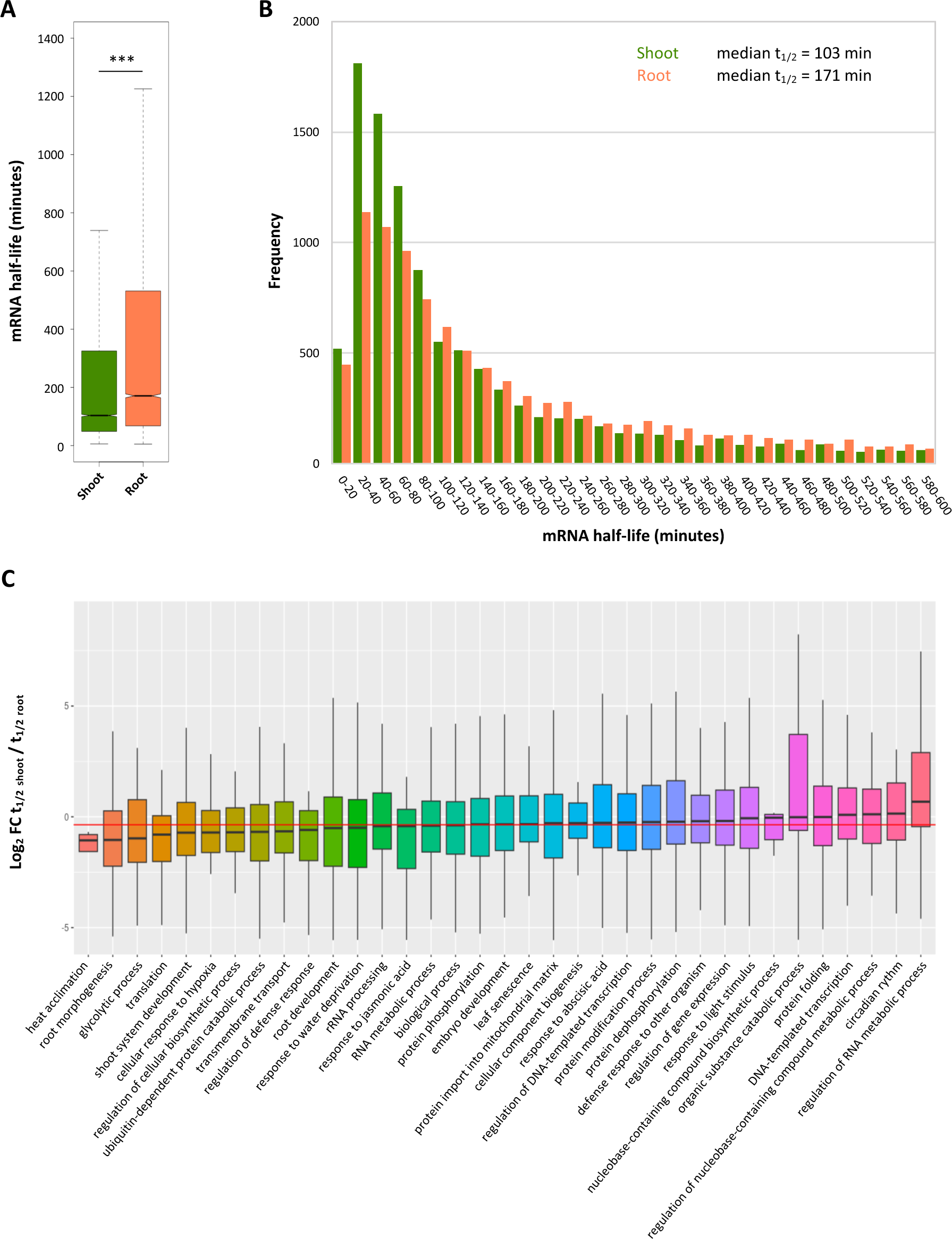
Analysis of mRNAs stability in WT shoot and root. A. Boxplots of mRNA half-lives in shoot and root (N= 12 306). B. Distribution of mRNA half-lives in shoot (green bars) and root (red bars). C. Distribution of the fold change between t_1/2_ shoot and t_1/2_ root separated into GO slim groups. The median value of the whole dataset is represented by a red line. Only groups with at least 10 members were used for the analysis.

### Contribution of CTRD in the general 5’-3’ mRNA turnover

The contribution of XRN4 in CTRD and in the general 5’-3’ mRNA turnover is presented in Figure 5. In shoot, the median half-lives in *xrn4* and XRN4ΔCTRD lines are similar (146 and 147 minutes respectively) and significantly higher than in WT (Figure 5A, Supplemental Table 2) indicating that CTRD contributes to decay of many mRNAs and supporting the idea of an important contribution in 5’-3’ mRNA decay in this tissue. In root, the median half-life is also significantly higher in *xrn4* (218 minutes compared to 171 minutes in WT) but surprisingly, the median half-life in XRN4ΔCTRD line is significantly lower (112 minutes versus 171 minutes in WT). The majority of decay rates are faster in XRN4ΔCTRD (Figure 5B, Supplemental Table 2). To exclude a transgenic line effect, we repeated mRNA half-life determination on the two independent XRN4ΔCTRD lines. The same phenomenon is observed with the two lines, supporting a biological effect due to the absence of CTRD in root (Supplemental Figure 9).

**Figure 5.**
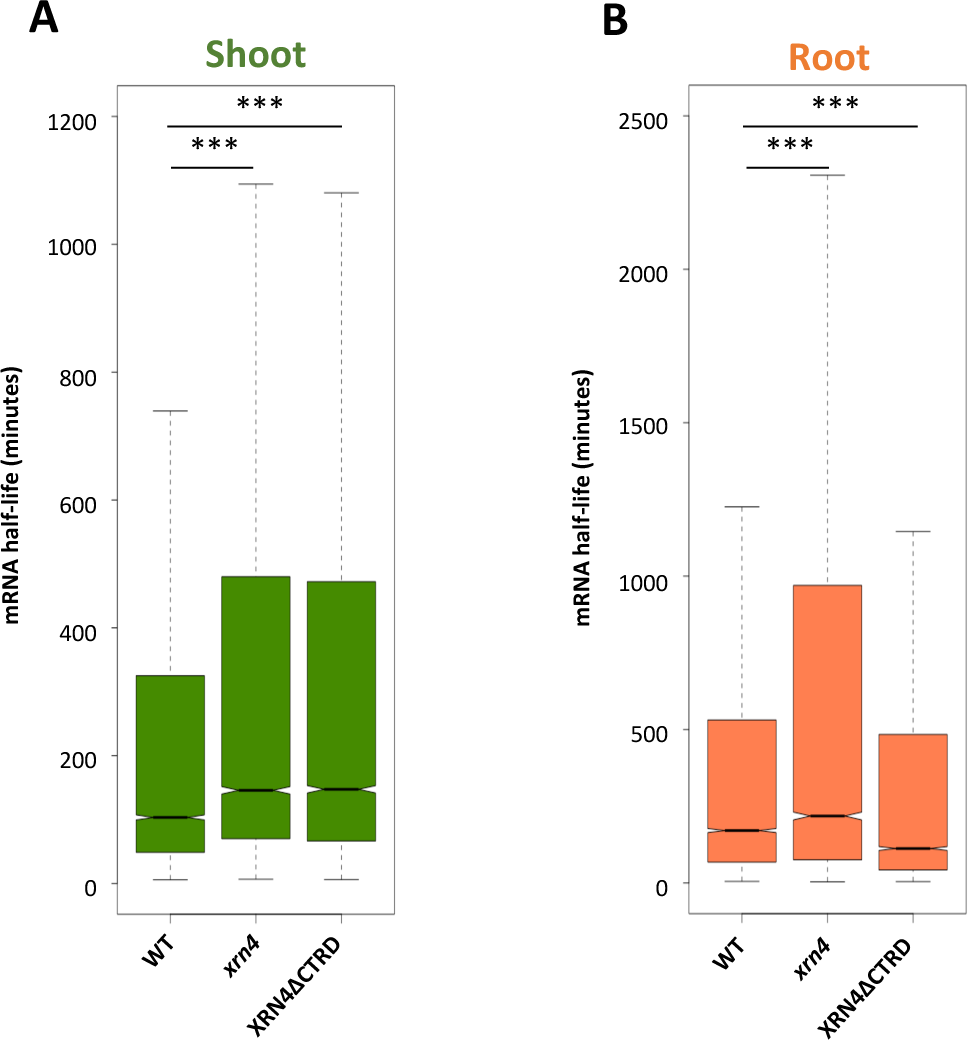
RNA half-life distributions in *xrn4* and XRN4ΔCTRD lines. Data are presented in shoot (A) or in root (B). A Wilcoxon test was performed to test significance. N= 12 306.

To determine the contribution of the two decay pathways for each mRNA in root and shoot, we used the likelihood function available in the RNAdecay package to assign a statistical genotypic effect to each transcript (13, 32). Modelling considered 5 combinations (called α groups) of genotypic effects (Supplemental Figure 10). An mRNA might show a different half-life in each genotype (α group 1) or show a similar half-life in two genotypes (α groups 2, 3 and 4), or show the same half-life in all genotypes (α group 5). In α groups 1 to 4, different subgroups exist based on the effects of *xrn4* and XRN4ΔCTRD lines on mRNA half-life (Supplemental Figure 10). Interestingly, the distribution of mRNAs in α groups is different between root and shoot suggesting distinct decay mechanisms in each organ. The α group 5.1 is the most abundant in shoot and root with 3 332 and 2 679 transcripts respectively, representing 27% (3 332/12 306) and 22% (2 679/12 306) of the population. These transcripts are not affected in *xrn4* and XRN4ΔCTRD lines (Supplemental Figure 10).

According to *xrn4* and XRN4ΔCTRD lines genotypic effects, we categorized subgroups in two main groups, ‘XRN4 targets’ and ‘RNA decay feedback targets’. We considered that the XRN4 target group consists of transcripts with slower decay (*e.g.* longer t_1/2_) in the *xrn4* and XRN4ΔCTRD lines than in WT (subgroups 1.4, 1.6, and 2.2) or with slower decay in *xrn4* and not affected in the XRN4ΔCTRD line (subgroup 4.2) (Supplemental Figure 10, Supplemental Table 2). The RNA decay feedback group encompasses transcripts with faster decay in *xrn4* or XRN4ΔCTRD lines compared to WT (subgroups 1.1, 1.2, 1.3, 1.5, 2.1, 3.1 and 4.1). Finally, since the biological relevance of the subgroup 3.2 (t_1/2_ XRN4ΔCTRD > t_1/2_ *xrn4* = t_1/2_ WT) was difficult to interpret, its transcripts were removed as well as those from subgroup 5.1 (not affected in the *xrn4* or XRN4ΔCTRD lines). We hereby retained 7 882 and 9 291 transcripts from shoot and root respectively, for further analyses. The distribution of the ‘XRN4’ and ‘RNA decay feedback’ target groups is presented in Figure 6A-B. Strikingly, the proportions of the two groups are totally different between organs, with 71% (5 634) of XRN4 targets and 29% (2 248) of RNA decay feedback in shoot compared to 39% (3 648) and 61% (5 643) in root, here again suggesting distinct decay regulations between shoot and root.

**Figure 6.**
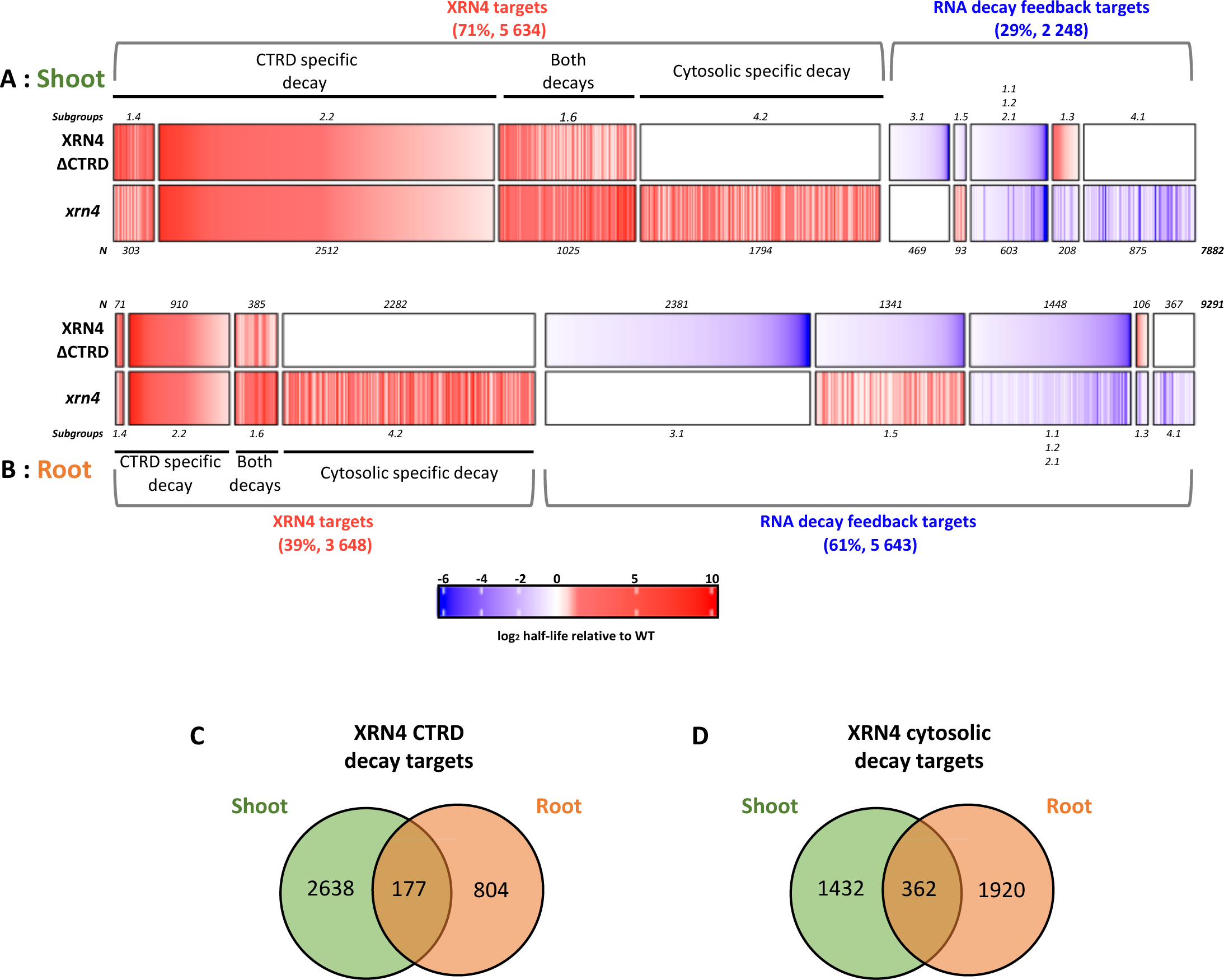
Identification of XRN4 co-translational and cytosolic targets and RNA decay feedback in shoot and root. Heat map representing t_1/2_ in XRN4ΔCTRD and *xrn4* relative to t_1/2_ in WT in shoot (A) or in root (B). Transcripts are organized in α subgroups (indicated on each cluster) and ordered in ‘XRN4 targets’ and ‘RNA decay feedback targets’ groups. N= 7 882 in shoot and 9 291 in root. C. Comparison of XRN4 CTRD specific targets (subgroups 1.4 and 2.2) in shoot and root. D. Comparison of XRN4 cytosolic specific targets (subgroup 4.2) in shoot and root.

To identify transcripts exclusively targeted by cytosolic or CTRD pathways, we reasoned as follows: 1-A transcript only targeted by CTRD pathway will have a similar increased half-life in both XRN4ΔCTRD and *xrn4* lines (subgroup 2.2). As increased half-life in XRN4ΔCTRD line is slightly higher than *xrn4* in subgroup 1.4, these transcripts were also considered as CTRD specific targets, 2-A transcript only targeted by the cytosolic pathway will have an increased half-life in *xrn4* and a half-life not affected in XRN4ΔCTRD line (subgroup 4.2), 3-A transcript targeted by both pathways will have an increased half-life in both XRN4ΔCTRD and *xrn4* lines but at lower amplitude in XRN4ΔCTRD line (subgroup 1.6).

In shoot, 32% of XRN4 targets (1 794 out of 5 634 XRN4 targets, subgroup 4.2) are only affected in *xrn4* and not in XRN4ΔCTRD line, suggesting that these transcripts are targeted only by XRN4 cytosolic decay. *A contrario*, 50% are exclusively targeted by the CTRD pathway (2 815, subgroups 2.2 and 1.4) while 18% (1 025) are targeted by both pathways (subgroup 1.6). These data are consistent with the t_1/2_ distribution observed in *xrn4* and XRN4ΔCTRD lines in shoot (Figure 5A) and suggest that CTRD is the major 5’-3’ mRNA decay pathway in shoot.

In root the situation is more complex. XRN4 targets represent only 39% of the population (3 648/9 291) with most of them targeted by cytosolic decay (2 282/3 648, subgroup 4.2). On the other hand, the majority of transcripts (61%, 5 643/9 291) presents a faster decay in *xrn4* and/or XRN4ΔCTRD line (Figure 6B). This phenomenon was already observed in the *Arabidopsis thaliana sov* mutant (13) and suggests that an RNA decay feedback mechanism takes place in the absence of cytosolic and/or CTRD decays. The most important effect is in subgroup 3.1 (2 381). Indeed, transcripts of this group are decayed faster only in XRN4ΔCTRD line. This subgroup was only represented by 469 transcripts in shoot. Interestingly, subgroups in which transcripts decay is faster in XRN4ΔCTRD line (1.1, 1.2, 1.5, 2.1, 3.1) are the only subgroups that increase in number between shoot and root suggesting that a specific feedback mechanism takes place in the absence of CTRD in root.

Next, we compared CTRD- and cytosolic-specific targets between organs and observed only a very limited overlap, suggesting that identical mRNAs are turned over by distinct means in different organs (Figure 6C-D).

In order to determine biological processes targeted by 5’-3’ mRNA decay in root and shoot, a Gene Ontology (GO) analysis was performed on XRN4 CTRD specific targets (subgroups 1.4 and 2.2), cytosolic specific targets (subgroup 4.2) and RNA decay feedback targets (subgroups 1.1, 1.2, 1.3, 1.5, 2.1, 3.1 and 4.1) (Figure 7). In both organs, XRN4 CTRD and cytosolic specific targets are part of very distinct biological processes. In shoot, mRNAs coding for various RNA processes such as ribosome biogenesis or ribonucleoprotein complex biogenesis are targeted by CTRD while mRNAs coding for vesicle localization, phosphate starvation or alpha-amino acid metabolic process are targeted by cytosolic decay (Figure 7).

**Figure 7.**
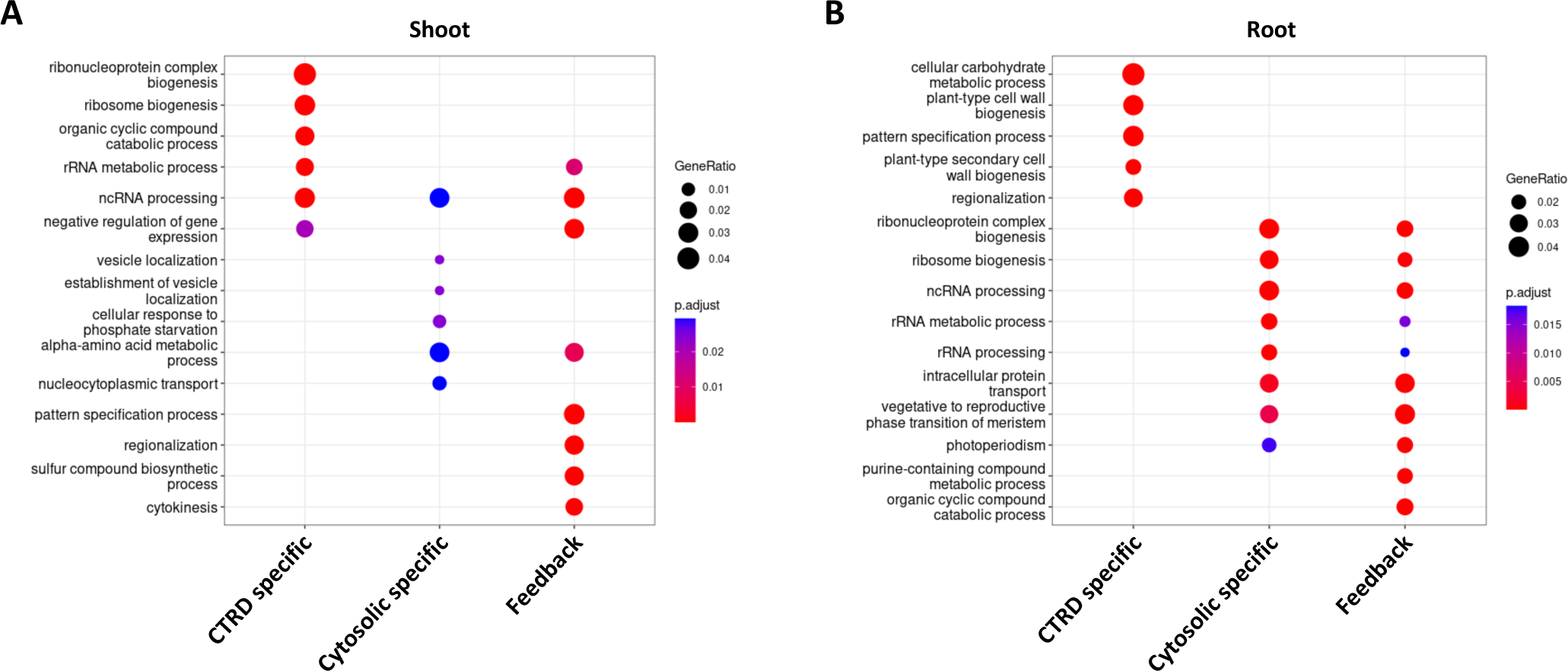
GO term analysis reveals specific biological functions targeted by cytosolic and co-translational decays. GO term analysis was performed on XRN4 CTRD specific targets (subgroups 1.4 and 2.2), XRN4 cytosolic specific targets (subgroup 4.2) and RNA decay feedback targets (subgroups 1.1, 1.2, 1.3, 1.5, 2.1, 3.1 and 4.1) in shoot (A) and root (B).

### Part of the feedback mechanism is mediated by the XRN4 cytosolic decay pathway

Many studies report the existence of a buffering system in which defects in turnover are compensated by a transcriptional readjustment, to maintain even steady-state mRNA levels (13, 39–41). To assess whether the faster decay rates observed in XRN4ΔCTRD line in root induce transcriptional feedback, we compared RNA abundances at T_0_ in XRN4ΔCTRD line and WT (Supplemental Figure 11). Using the whole dataset, no clear differences are observed between XRN4ΔCTRD line and WT. By contrast, CTRD specific targets present higher abundance in XRN4ΔCTRD line while RNA decay feedback targets present lower abundance suggesting that the loss of CTRD does not lead to transcriptional feedback (Supplemental Figure 11). Faster decay rates observed in the root of the XRN4ΔCTRD line suggest that an alternative decay mechanism takes place in the absence of CTRD. To test the possibility of a feedback mechanism mediated by the XRN4 cytosolic pathway, we created an additional XRN4ΔCTRD line but impaired in its 5’-3’ exonucleolytic activity. To do so, we performed a protein sequence alignment between AtXRN4 and diverse XRN orthologues (Supplemental Figure 12A). In *Drosophila melanogaster*, it was reported that the substitution of arginines R100 and R101 into alanines abolishes DmXRN1 activity. These two arginines are conserved and present in AtXRN4 at positions 118 and 119. Thus, we created the *AtXRN4ΔCTRD(R_118_A/R_119_A)* allele, that we expressed under the control of its own promoter in the *xrn4* background. We selected a transformant line expressing transgene levels similar to that of endogenous XRN4 and transgenic XRN4ΔCTRD (Supplemental Figure 12B). We next measured half-lives of candidate transcripts, through ddPCR-monitoring of RNA levels following cordycepin treatment (33), in wild-type, *xrn4*, XRN4ΔCTRD and XRN4ΔCTD(R_118_A/R_119_A) backgrounds (Figure 8 and Supplemental Figure 13). First, to confirm that the XRN4ΔCTRD allele retains normal and similar catalytic activities both in root and shoot, we selected three transcripts amongst the list of those identified as only targeted by the XRN4 cytosolic decay in both organs (subgroup 4.2, Figure 6A-B). For all three of them, mRNA half-life measurement revealed similar half-lives in WT and XRN4ΔCTRD lines in both shoot and root while, their half-lives significantly increased at the same amplitude in *xrn4* and XRN4ΔCTRD(R_118_A/R_119_A) lines (Figure 8A, Supplemental Figure 13A). This first analysis confirms that insertion of the mEGFP into AtXRN4 loop L3, does not affect its catalytic activity, as is the case for ScXRN1. It also comes as a validation of the transcriptome-wide assays that identified the mRNAs from subgroup 4.2 as cytosolic XRN4 targets. Next, we selected three transcripts only targeted by CTRD in both shoot and root (subgroup 2.2, Figure 6A-B). mRNA half-life measurement revealed significantly higher mRNA stability in *xrn4*, XRN4ΔCTRD and XRN4ΔCTRD(R_118_A/R_119_A) lines compared to WT in both organs (Figure 8B, Supplemental Figure 13B) supporting that transcripts from subgroups 2.2 are CTRD XRN4 targets. Finally, we selected transcripts from subgroup 1.5 to validate and analyse the feedback mechanism observed in root (Figure 8C, Supplemental Figure 13C). Interestingly for transcripts of subgroup 1.5, the feedback mechanism observed in XRN4ΔCTRD line is totally abolished in XRN4ΔCTRD(R_118_A/R_119_A) line suggesting that XRN4 cytosolic decay can compensate partially the absence of CTRD in root (Figure 6B).

**Figure 8.**
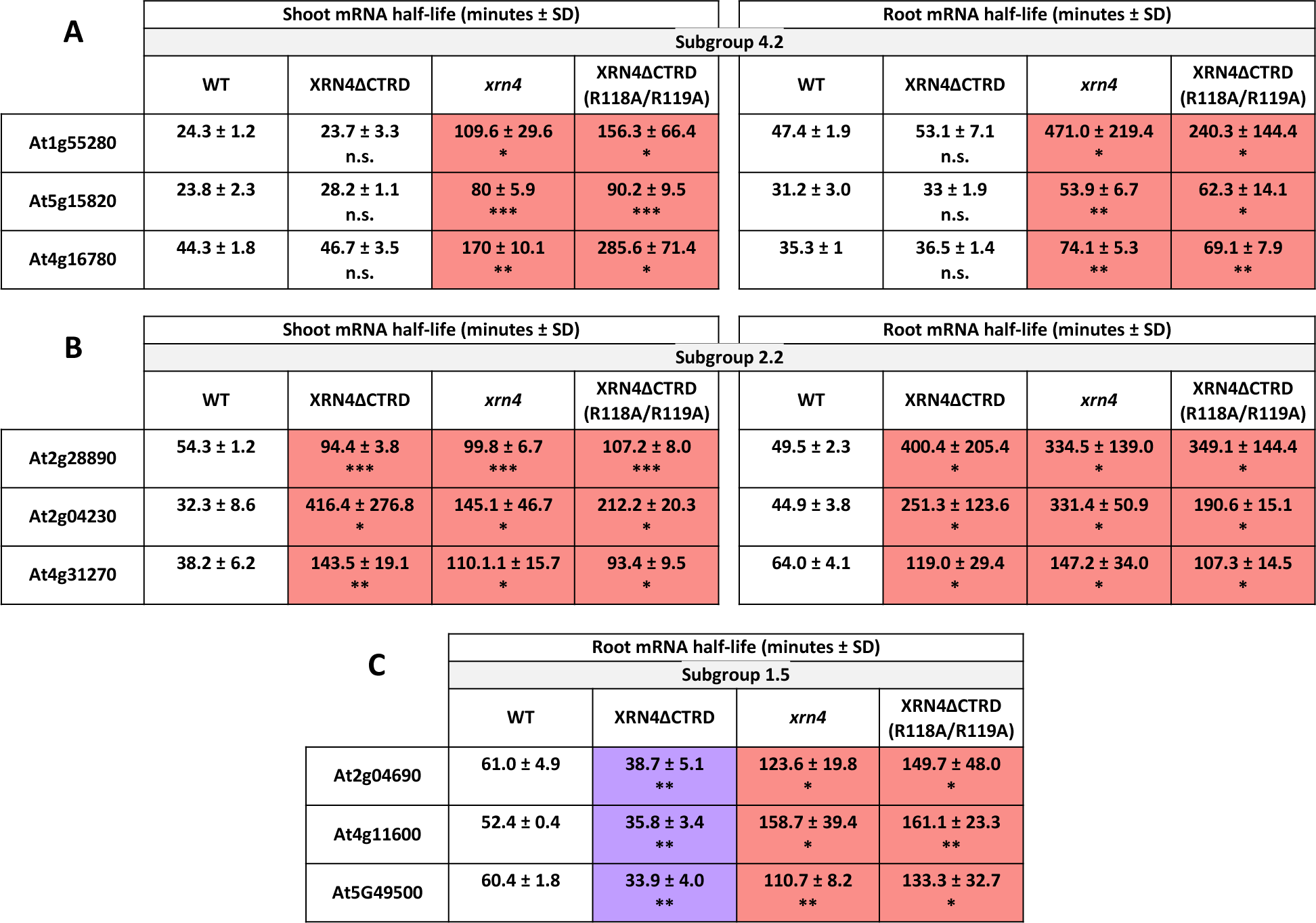
Part of the feedback mechanism is mediated by the XRN4 cytosolic decay pathway. mRNA half-life was determined according to data presented on Supplemental Figure 13. mRNA stability was determined *in vivo* after cordycepin treatment followed by RT-ddPCR on selected transcripts from subgroups 4.2 (A), 2.2 (B) and 1.5 (C). N= 3 biological replicates. Normalization was performed using Luciferase RNA spike-in. Mean ± SD. The significance was tested against WT and is indicated below each mRNA half-life. The color chart indicates slower or faster decay as on Figure 5.

### XRN4ΔCTRD line presents contrasting phenotypes in shoot and root

As mRNAs half-life distribution in *xrn4* and XRN4ΔCTRD lines reveals similar profiles in shoot and opposite profiles in root (Figure 5), we tested if this discrepancy could be associated with different phenotypes at the organ level. Several phenotypes were already observed in *xrn4*, such as defects in root growth under various conditions (6, 8) or leaf development (9). We analyzed these different phenotypes in XRN4ΔCTRD line in comparison to WT, *xrn4* and a complemented XRN4-GFP line (Figure 9). At leaf level, XRN4ΔCTRD line presents phenotypes similar to that of *xrn4* such as serrated leaves, higher leaf fresh weight and greater number of leaves (Figure 9A-C). However, at the root level, XRN4ΔCTRD line presents phenotypes opposite to that of *xrn4* (Figure 9D-E). Primary roots are longer in XRN4ΔCTRD line compared to WT while *xrn4* presents a shorter primary root (Figure 9D). Interestingly, this phenomenon is still observed under stress conditions such as salt stress (Figure 9E). To correlate these phenotypes to transcripts stability, we focused on transcripts associated with the GO terms “root morphology” and “root development”. Interestingly, among them, 111 transcripts present distinct mRNA half-lives between *xrn4* and XRN4ΔCTRD line: 41 transcripts have opposite half-lives in *xrn4* and XRN4ΔCTRD line (subgroup 1.5 or subgroup 1.3) and 70 present faster decay rates in XRN4ΔCTRD line and are not affected in *xrn4* (Supplemental Table 4). These data could suggest that the misregulation of mRNA stability for transcripts involved in root morphogenesis/development could affect root growth. All together, these phenotypic analyses reveal a strong correlation between decay rates alteration and organ phenotype.

**Figure 9.**
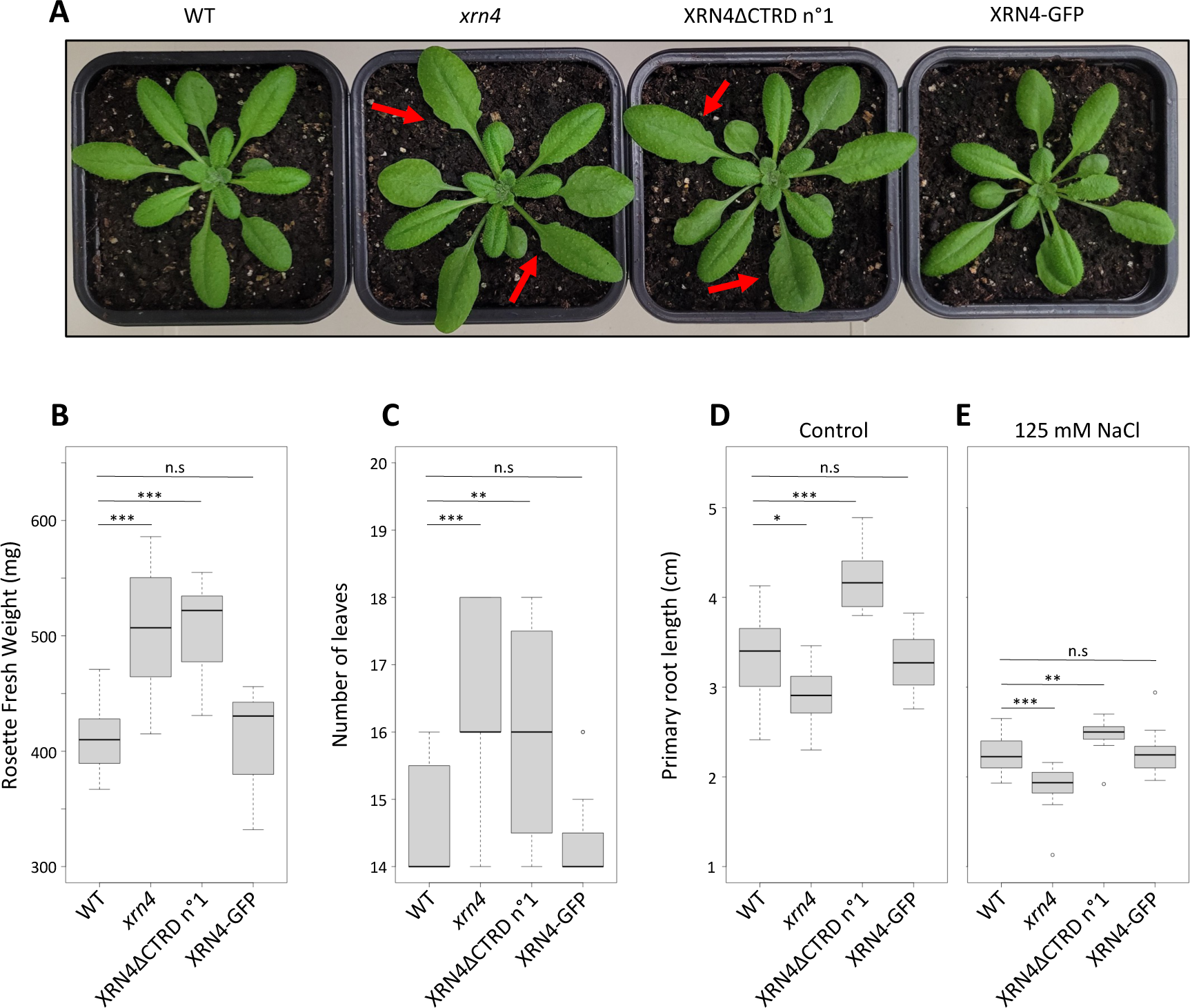
XRN4ΔCTRD line presents similar phenotypes as *xrn4* at leaf level and opposite phenotypes at root level. WT, *xrn4*, XRN4ΔCTRD line, and XRN4-GFP complemented line rosette morphology (A), rosette fresh weight (B), number of leaves (C), primary root length under normal (D) and salt stress conditions (E). Red arrows indicate serrated leaves. A t.test was performed to test significance. N> 18.

## DISCUSSION

mRNA decay plays an important role in the post-transcriptional regulation of gene expression. In eukaryotes, two major pathways can together turn over the transcriptome, the 5’-3’ mRNA decay and the 3’-5’ mRNA decay (10). In plants, defects in 5’-3’ mRNA decay can induce drastic developmental and stress response phenotypes revealing the importance of this pathway (3, 4, 6, 7, 9, 42). The general 5’-3’ mRNA decay was placed in the cytosol after ribosomes release.

However, for many years, it has also been proposed that mRNA decay can occur on translated mRNAs through a 5’-3’ co-translational mRNA decay mechanism (15, 16, 20). While players, enzymes and cofactors of the 5’-3’ turnover are well recognized, the importance either cellular or physiological and the relative contribution of each pathway has never been assessed in any organism. As a matter of fact, uncoupling the 5’-3’ decay pathways proved difficult because they are catalysed by the same enzymes, the main one being XRN4 in *Arabidopsis thaliana*. Here, we developed a transgenic line impaired in CTRD that allows the discrimination between 5’-3’ cytosolic and 5’-3’ CTRD pathways. Using a genome-wide mRNA decay analysis, we determined the importance of CTRD in both shoot and root and provide evidence that it plays a crucial role in root development.

mRNA decay analysis in WT in shoot and root reveals diverse range of decay rates (from minutes to hours) with a respective median value of 103 and 171 minutes. This range is quite similar to previous mRNA decay analysis made in *Arabidopsis thaliana* (13, 43) but reveals an organ-specific pattern. This organ-specific mRNA stability was already observed in different organisms. As an example, in *Drosophila melanogaster*, stabilization of transcripts involved in cell-fate decisions and axonogenesis is an important process for neural development (44). In human, changes in mRNA stability contribute to fibroblast quiescence maintenance (45). By analyzing *cis*- and *trans*-elements, we found that SKI2 transcript is less stable in root compared to shoot. We also found that PAT mRNA decapping factors (PAT1, PAT1H1, PAT1H2) are less abundant in root compared to shoot. The SKI complex (composed of SKI2, SKI3 and SKI8) is involved in the unfolding of mRNAs to assist exosome-mediated decay (46). The SKI complex is important at the leaf level as together with uridylation, they regulate RNA degradation pathway important for photosynthesis (47). This is consistent with the extreme stability of SKI2 mRNA in our shoot half-life dataset. In *Arabidopsis thaliana*, PAT1 has been implicated in pathogen response where it activates decay of specific transcripts (48). More recently, it was proposed that PAT1H1, PAT1H2 harbor specific and overlapping functions with PAT1 during plant growth (49). However, their importance at the organ level was never tested. More investigations will be needed to better assess the importance of decay factors in mRNA stability at the organ level.

The importance of the 5’-3’ mRNA decay was revealed by the distribution of mRNA half-lives in *xrn4*. In *xrn4*, the median half-life was 43 and 77 minutes higher than WT in shoot and root respectively indicating that XRN4 contributes to decay of many mRNAs. These data are in agreement with the contribution of the decapping (through VCS) in the decay of the transcriptome where 68% of transcripts present a VCS-dependent decay (13). The importance of CTRD was revealed by the distribution of mRNA half-lives in the XRN4ΔCTRD line. In shoot, the contribution is clear with similar half-life distributions in *xrn4* and XRN4ΔCTRD lines (Figure 5A). The α group pattern in shoot reveals that most of the 5’-3’ degradation is mediated by CTRD (Figure 6A). Interestingly, only few transcripts are targeted by both CTRD and cytosolic decays (α group 1.6) meaning that transcripts are mostly targeted by one of the two pathways.

The specific inactivation of the XRN4-mediated CTRD in root results for a large fraction of the root and to a lesser extent of the shoot transcriptomes, in a counter-intuitive phenomenon where thousands of transcripts decay faster (Figure 5B and Figure 6B). Such phenomenon was previously reported in yeast and *Arabidopsis thaliana* and recognized as a buffering process, of RNA decay defects (40, 41, 50). In yeast, the reduction of decay rates induces the reduction of the transcription rates to maintain mRNA steady-state at proper levels (41, 50). This mechanism was called RNA buffering and is mediated by the nucleocytoplasmic shuttling of ScXRN1 (41). In fact, two NLS sequences were recently identified in ScXRN1 that are important for its import to the nucleus and for transcription rate modulation (39). In plants, no direct ortholog of XRN1 exists. *Arabidopsis thaliana* indeed carries three XRNs, AtXRN2, AtXRN3 and AtXRN4 that are structurally similar to ScXRN2/RAT1 (51). The deletion of a bipartite NLS during duplication may have given rise to an AtXRN4 cytoplasmic exoribonuclease that have cytoplasmic functions similar to that of ScXRN1 (51). Thus, AtXRN4 is probably not able to perform nucleocytoplasmic shuttling explaining the absence of transcriptional feedback in the XRN4ΔCTRD line (Supplemental Figure 11).

In *xrn4* and XRN4ΔCTRD lines, the 3-nt periodicity and 5’P reads accumulation -16/-17 nt before stop codons are not totally abolished (Figure 3A). *A contrario*, in *Saccharomyces cerevisiae,* deletion of XRN1 totally abolishes this periodicity and 5’P reads accumulation before stop codons (15). In addition, the 3-nt periodicity is less pronounced in *Arabidopsis thaliana* and *Schizosaccharomyces pombe* compared to *Saccharomyces cerevisiae* (15, 22, 52). This could probably explain why the 3-nt periodicity seems still present in *xrn4* and XRN4ΔCTRD lines (Figure 3A, (16)), suggesting distinct co-translational decay mechanisms between organisms.

Faster decay rates in root of XRN4ΔCTRD line suggest that another decay mechanisms takes place in the absence of CTRD. Using XRN4ΔCTRD(R_118_A/R_119_A) line, we demonstrated that part of this feedback is mediated by the XRN4 cytosolic decay (subgroup 1.5, Figure 8C). However, transcripts from other subgroups present also RNA decay feedback. We and others proposed that XRN4 is not the only exoribonuclease involved in CTRD (16, 53). Interestingly, in *fry1* mutant, 5’-3’ mRNA decay seems more repressed than in *xrn4* (53). In *fry1* mutant, a constitutive accumulation of 3’-phosphoadenpsione 5’-phosphate (PAP) is observed resulting in inhibition of XRN activity (54). These data suggest that in addition to XRN4, another exoribonuclease sensitive to PAP, could be involved in CTRD. Whether the feedback mechanism is mediated by another co-translational decay pathway is still an open question.

Why and how at the molecular level, the compensation phenomenon to the loss of CTRD is significantly more important in root than in shoot is another interesting question that emerges from our work and that will need further studies to answer. However, consistent with the differences in compensation and the distinct contributions of the CTRD to overall 5’-3’ mRNA turnover, we report that the CTRD takes on distinct physiological importance in root and shoot (Figure 9). Indeed, the phenotypes observed in the XRN4ΔCTRD line correlate with alterations of decay rates. In shoot, the *xrn4* and XRN4ΔCTRD lines present similar alterations of decay rate resulting in similar phenotypes while in root, opposite alterations of decay rates are observed resulting in opposite phenotypes. Taken together, our data report that CTRD plays important roles in the fine-tuning of mRNA turnover and is essential for proper plant development. The development of this XRN4ΔCTRD line will give to the community the opportunity to analyze deeper the importance of CTRD in plant development and stress response.

Overall, our data highlight the importance of CTRD in mRNA decay and more largely in mRNA metabolism. As coupled to translation, this pathway can significantly affect translation efficiency and protein production (16). As this pathway is conserved in many organisms, highly regulated by development and stress (7, 15, 16, 19–22, 25), its contribution to translation efficiency should be considered with more attention in future studies.

## DATA AVAILABILITY

The RNAseq and 5’Pseq datasets generated in this study have been deposited in the Short Read Archive with the accession code PRJNA1018408 and PRJNA1023397 respectively.

## SUPPLEMENTARY DATA

Supplementary Data are available at NAR online.

## ACKNOWLEDGEMENTS

This study is set within the framework of the “Laboratoires d’Excellences (LABEX)” TULIP (ANR-10-LABX-41) and of the “École Universitaire de Recherche (EUR)” TULIP-GS (ANR-18-EURE-0019). The authors gratefully acknowledge Jean-Marc Deragon for critical reading of the manuscript.

## FUNDING

Agence Nationale de la Recherche [ANR-21-CE20-0003 to R.M.]

Agence Nationale de la Recherche [ANR-17-CE20-0007 to C.B.A.]

